# Trapping the exotic weevil *Cnestus mutilatus* with Isopropanol

**DOI:** 10.1101/2023.11.26.568738

**Authors:** Teresa C. Justice, Michael J. Justice

## Abstract

The ambrosia beetle Cnestus mutilatus Blandford, 1894 (Coleoptera: Curculionidae: Scolytinae: Xyleborini) is native to Asia and is currently an exotic species in North America. This study tested whether this species can be trapped with isopropanol as bait. Lindgren multiple-funnel traps were deployed in the piedmont of south-central Virginia, USA. The experimental traps had isopropanol in the collection cups. For comparison, other traps had ethanol or soapy water. Both alcohols were effective for trapping C. mutilatus. More specimens were captured using ethanol. Isopropanol and ethanol may play similar roles in the chemical ecology of ambrosia beetles.

## INTRODUCTION

The ambrosia beetle *Cnestus mutilatus* Blandford, 1894 (Coleoptera: Curculionidae: Scolytinae: Xyleborini) is native to Asia but was detected in the southern United States in 1999 (1, 2). The species has since spread widely and quickly (3), utilizing a suite of adaptive traits: a wide range of host trees, haplodiploidy, inbreeding, sheltering in wood for most of their life cycle, sub-sociality, fungus gardening, and the ability to move from tropical to temperate climates (4–6).

It is possible that *C. mutilatus* will not become a pest in its new geographic range. Most ambrosia beetles take advantage of the weakened defenses of stressed or dying trees, in which they typically sow nonpathogenic fungi (7) that contribute to decomposition and nutrient recycling.

However, timber- and tree-associated industries need to remain wary of their destructive potential. The woody tissues of dying and dead trees ferment residual sugars, producing several short-chain alcohols such as ethanol (8, 9). A small proportion of these alcohols volatize (10) and attract the beetles, which will tunnel breeding galleries into the woody tissues. Stressed nursery trees and many wood products also volatize alcohols, and holes from any resulting beetle galleries reduce the structural integrity, esthetics, and market value of these products. Examples include damage to wine or beer casks (8), overwatered or freeze-stressed nursery stock (11), recently felled timber (12), and freshly sawn lumber (13–15). *Cnestus mutilatus* has even been documented boring holes into the plastic walls of ethanol-gasoline containers (16). Ambrosia beetles could also cause ecological damage by seeding the spread of microbes, including their primary ambrosia fungus, secondary fungi, nematodes (17), mites, and bacteria (18). Fungi transported by ambrosia beetles can dramatically alter the natural progression of wood decomposition (19, 20) and could be phytopathogenic (21, 22). *Geosmithia morbida* (Ascomycota), the fungal pathogen that causes Thousand Cankers Disease in walnut trees (*Juglans* spp.), is known to be carried by *C. mutilatus* (23, 24). Given this potential for both economic and ecological damage, *C. mutilatus* merit close monitoring (25, 26).

Traps for ambrosia beetles are typically baited with slow-release ethanol preparations. However, propan-2-ol (hereafter referred to as isopropanol), while being used in traps as a killing agent and preservative, seemed to attract *C. mutilatus* (personal observations). There would be considerable advantages to having an inexpensive and easy-to-obtain chemical serve all three functions of attractant, killing agent, and preservative without the need for slow-release mechanisms. The purpose of this study was to gather experimental evidence that isopropanol can be used as such in traps for *C. mutilatus*.

## METHODS

Trapping took place at four sites near Lynchburg, Virginia, USA. All trapping sites were in residential neighborhoods and proximal to water and small patches of deciduous forest. All traps were multiple-funnel traps hung so the bottom was 1-1.5 m above the ground. Inserted into each trap’s collection cup was a close-fitting glass jar containing either an alcohol or a control. The jar’s inner diameter was approximately 10cm, so the liquids were evaporating from a surface of about 79 cm^2^.

### Sites 1 and 2

On 02 Jan 2021, two 12-funnel Lindgren traps (27) were deployed: one at 37.37379°N, 79.20182°W (Site 1) and another at 37.37396°N, 79.20164°W (Site 2). These traps were fitted with extended rain shields and secured with rope tethers to limit swaying. Initially, 150 mL of 91% isopropanol was used; this was increased to 200 mL during the warmer months to compensate for an increased evaporation rate.

Immediately after the first *C. mutilatus* was captured, two control 12-funnel traps were deployed in the same way, each hung about 2 m from a trap with isopropanol. Control traps often use ethylene glycol or propylene glycol, but these molecules contain alcohol groups and were deemed inappropriate for comparison with alcohol baits. Instead, control traps used 150 mL distilled water with a surfactant (a small shaving of Dove brand fragrance-free bar soap). These two experimental and two control traps were inspected almost daily. Debris such as spider webs and fallen leaves was removed, and the isopropanol was topped up if needed. For one year, on every seventh day, specimens were collected and the alcohol or soapy water was replaced. The isopropanol traps were emptied and refilled more frequently when Green June Beetles (*Cotinis nitida* Linnaeus, 1758) and American Carrion Beetles (*Necrophila americana* Linnaeus, 1758) were caught in large numbers.

### Sites 3 and 4

Two very different arrangements of Lindgren traps were deployed at two additional sites from April to October 2021. These traps were inspected twice per week; specimens were collected and the fluids replaced at least once per week. One site (37.38589°N, 79.24818°W; Site 3) had isopropanol and control traps, as above, but these were four-funnel traps hung 16 m apart. This size and distance should greatly reduce the control traps’ interception of beetles in flight toward the isopropanol trap. The last site (37.38816°N, 79.25908°W; Site 4) had three four-funnel traps hung in a line and spaced at 1 m intervals. At either side (thus 2 m apart) was a trap baited with 91% isopropanol or 40% ethanol. The position of these two alcohol traps alternated weekly. The middle trap contained soapy water for control.

*Cnestus mutilatus* were identified using the key in Gomez *et al*. (28). All captures were of females, as this is the only sex that disperses from the galleries. Voucher specimens will be deposited with the Florida State Collection of Arthropods (contact: Paul Skelley). Weather data were obtained from the U.S. National Weather Service (weather.gov/wrh/climate?wfo=rnk) and Weather Underground (wunderground.com). Double sine-wave integrals for Degree-Days were calculated using the weather data and Excel, and checked against the computations provided by two online services (uspest.org/dd/model_app and ipm.ucanr.edu/WEATHER/index.html). Derived variables and statistics were calculated using Excel.

## RESULTS

### Baits and Captures

Captures are summarized in Table 1. The first capture was on April 11 at Site 1; at the time, there was only an isopropanol trap deployed there (see Methods), so this capture was not influenced by the presence of another trap or other *C. mutilatus* in this trap. At Sites 1, 2, and 3, which simply compared isopropanol to soapy water control, capture numbers were significantly higher in isopropanol (binomial tests, each *p* value < 0.0001). Hundreds were captured in isopropanol versus a single specimen in soapy water; it is very unlikely that control traps were regularly intercepting beetles in flight on their way to an isopropanol trap.

**TABLE 1.** Captures of *C. mutilatus* by location and bait. Sites 1, 2, and 3 compared Isopropanol to Soapy Water. Site 4 included a trap with Ethanol for comparison.

|  | Isopropanol | Soapy Water | Ethanol | Total |
| --- | --- | --- | --- | --- |
| Site 1 | 440 | 0 | N/A | 440 |
| Site 2 | 116 | 1 | N/A | 117 |
| Site 3 | 165 | 0 | N/A | 165 |
| Site 4 | 121 | 6 | 513 | 640 |
| Total | 842 | 7 | 513 | 1362 |

Site 4 had an ethanol-baited trap in addition to isopropanol and control. Captures were significantly higher in ethanol compared to isopropanol (binomial *p* < 0.0001). Capturing only six specimens in the control trap, which was hung between the isopropanol and the ethanol traps at this site, strongly suggests the traps were not regularly intercepting beetles in flight toward another trap. Using parametric least squares regression, the weekly number of captures in one alcohol can be predicted from the captures in the other alcohol (*F*_1, 24_ = 12.62, *r*^2^ = 0.34, *p* = 0.0016; Ethanol = 3.40*Isopropanol + 3.90; Isopropanol = 0.10*Ethanol + 2.65).

The first captures occurred after only several Degree-Hours of air temperature above 27 °C had accumulated (Table 2). There were three distinct peaks in the captures: at the end of April, end of May, and mid-June (Figure 1). The peaks and troughs for isopropanol and ethanol coincided in time but were smaller in magnitude for the isopropanol traps.

**TABLE 2.**
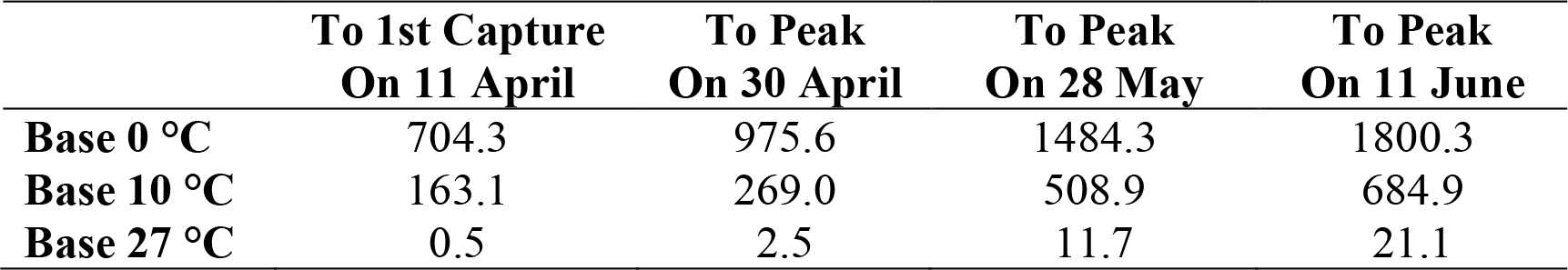
Celsius Degree-Days accumulated prior to key dates in 2021. The biofix date is 01 January 2021. Calculations used the double sine wave method. Base 27 °C is included because it appears to be a threshold for the appearance of *C. mutilatus* in traps. Base 0 °C and Base 10 °C are included because they are popular for Degree-Day calculations and could be useful for predictions. Note that heat input would not be for development because this species overwinters as adults (cf. 29).

|  | To 1st Capture<br>On 11 April | To Peak<br>On 30 April | To Peak<br>On 28 May | To Peak<br>On 11 June |
| --- | --- | --- | --- | --- |
| **Base 0 °C** | 704.3 | 975.6 | 1484.3 | 1800.3 |
| **Base 10 °C** | 163.1 | 269.0 | 508.9 | 684.9 |
| **Base 27 °C** | 0.5 | 2.5 | 11.7 | 21.1 |

**FIGURE 1.**
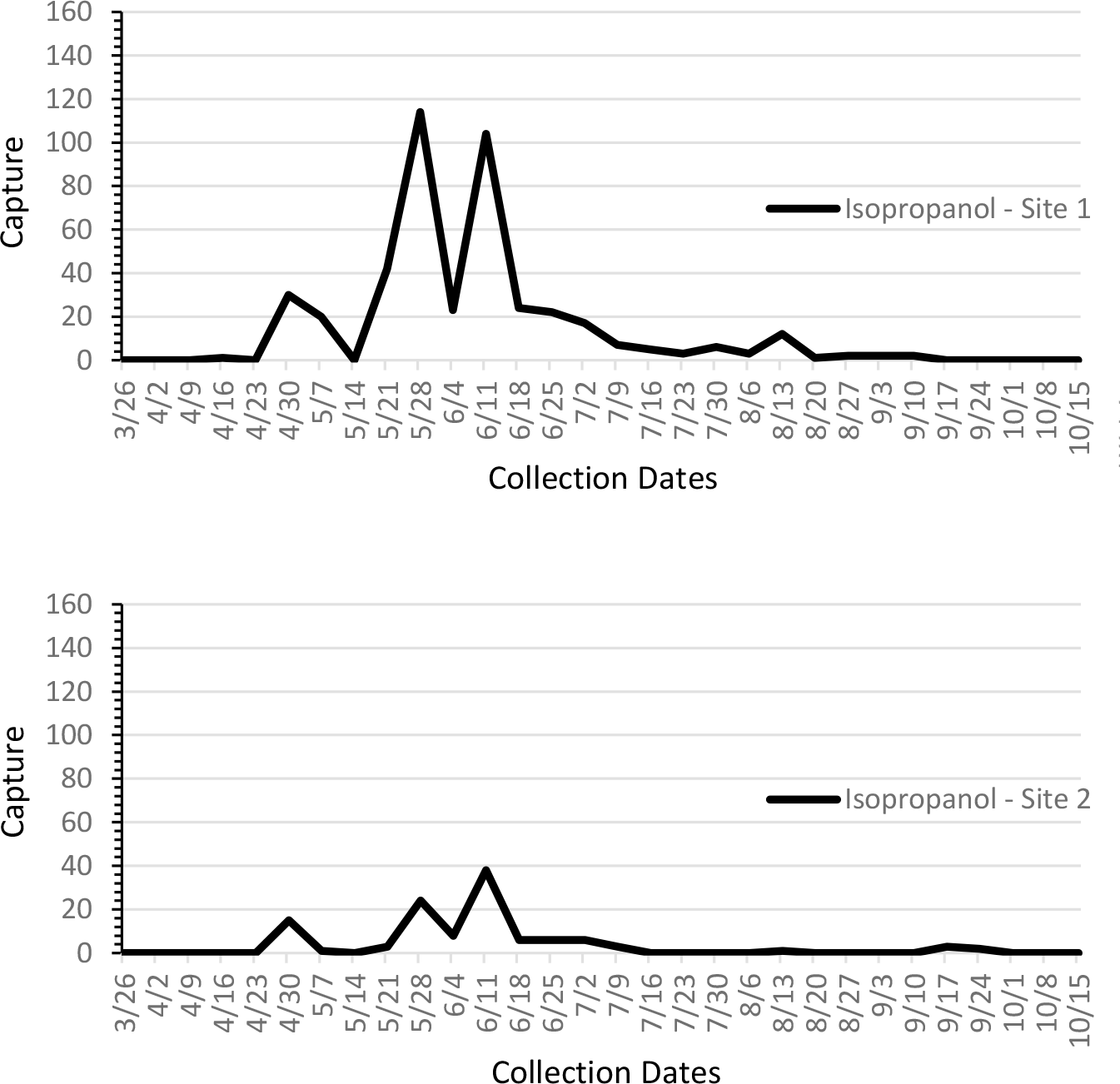
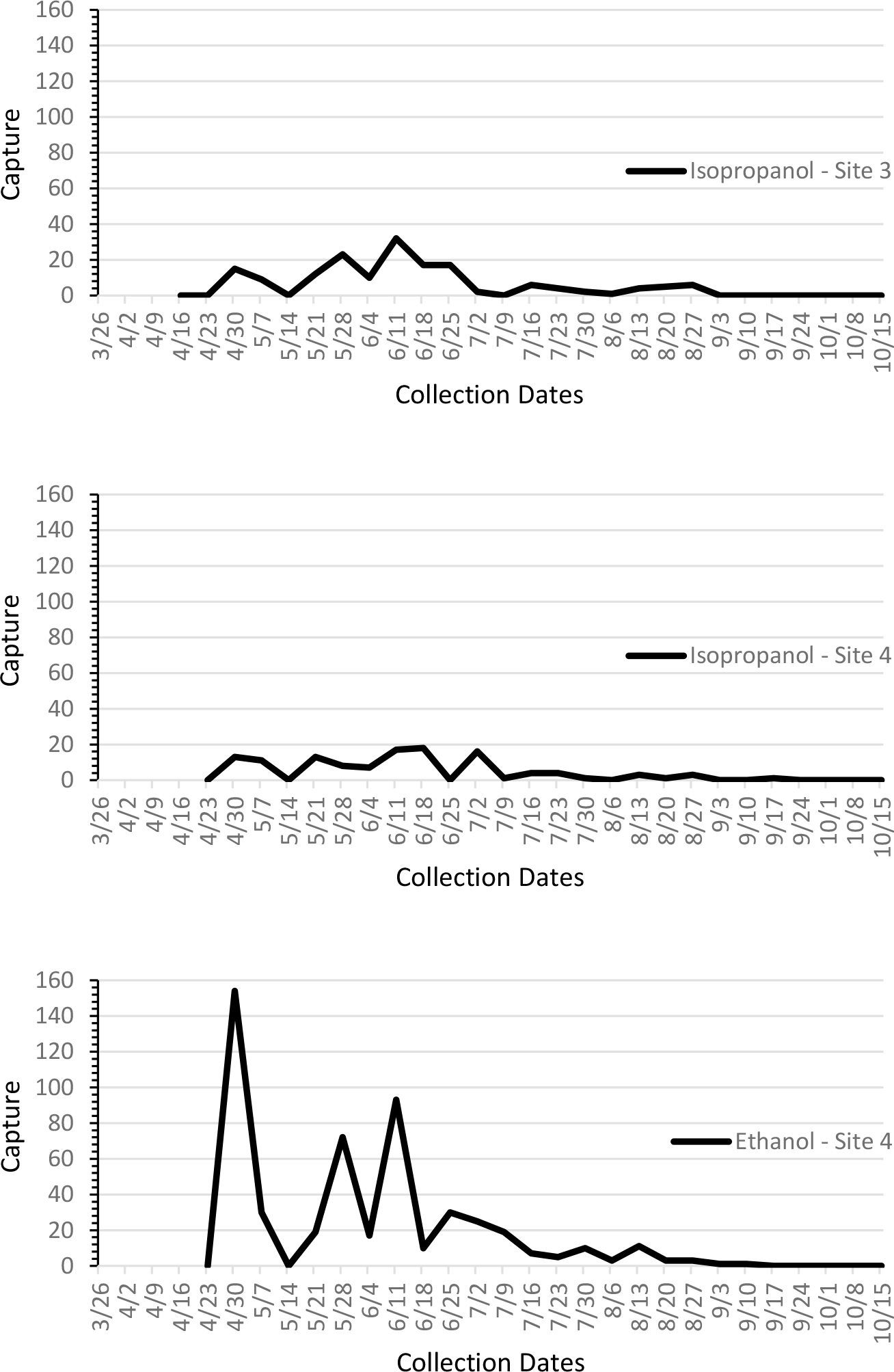
Weekly capture data. The *x*-axis tick marks are the collection dates in month/day format.

### Correlations Between Captures and Weather

Correlation coefficients were calculated using weekly data from the *n* = 24 weeks of captures between April and October 2021 (with *df* = 22, the Critical Value of *r* at α = 0.05 is ±0.4044). “Weekly total captures” was calculated as the sum of all captures in both alcohols at all four sites. Three different temperature variables were calculated by averaging (over the week) the daily maximum, minimum, and average temperatures. As measured, temperature did not correlate with the number of captures: using maximum temperatures, *r* = +0.12; minimum temperatures *r* = -0.09, and average temperatures *r* = +0.02.

Other weather variables were chosen based on the likelihood they might affect either the beetles’ activities or the growth of their ambrosia fungi. Of these, the only variable that significantly correlated with captures was the total amount of rainfall accumulated over the week prior to the capture count. (*r* = +0.55, two-tailed *p* < 0.01). None of the following variables was significantly correlated with captures: maximum relative humidity (*r* = +0.16), minimum relative humidity (*r* = -0.13), average relative humidity (*r* = +0.02), average wind speed (*r* = +0.19), maximum barometric pressure (*r* = +0.12), minimum barometric pressure (*r* = -0.07), and average barometric pressure (*r* = +0.10).

## DISCUSSION

The data indicate that *C. mutilatus* can be trapped with isopropanol. The pattern of weekly capture numbers was very similar for isopropanol and ethanol, but the number caught with isopropanol was consistently smaller. Captures in one alcohol can fairly accurately predict the number of captures in the other.

Insofar as capture data can be used to infer emergence and flights (see 30), *C. mutilatus* appears in late April in large numbers, possibly in response to temperatures remaining above 27 °C for several hours (cf. 31–33). The initial peak is followed by a trough in mid-May and another, larger peak (or possibly two) about 30–40 days later in late May to mid June. The separation of these peaks could reflect varying latencies in the response to springtime conditions (32), but they are separated by enough time to also be the result of separate broods if the species is bivoltine in this region (see 34 for phenology information). For comparison, two peaks were also seen in Mississippi (35) and Georgia (36) using Lindgren traps, but only a single peak was seen in Kentucky using Baker bottle traps (37).

Isopropanol is known to be bioactive in several contexts. In addition to attracting various insects into traps (38–46), it is known to volatize from insects (47), stimulate oviposition (48), and evoke antennal electrophysiological responses (46, 47). Isopropanol also volatizes from some plants (38, 41, 48–54) and fungi (55, 56). Stress-related processes such as fermentation (57, 58) and amino acid metabolism (59–61) can result in the production of isopropanol. Thus, the potential exists that isopropanol, like ethanol, might be volatized by trees under stress and used by *C. mutilatus* to locate such trees for their galleries.

Once in the galleries, bark beetles cultivate ambrosia fungi for food. Ethanol volatizing from the stressed woody tissues is toxic to fungi. The ambrosia fungi survive by using alcohol dehydrogenase enzymes (ADHs) to bind and detoxify the ambient ethanol (62, 63). Other fungi in competition with the ambrosia fungi are culled by the ambient ethanol. Isopropanol is similarly volatile, toxic (64), and able to affect fungal growth (65, 66). Many ADHs that bind ethanol will also bind isopropanol, and some ADHs have high specificity to isopropanol (60, 67, 68). Thus, it is possible that isopropanol could play similar roles to ethanol in the galleries of ambrosia beetles.

Isopropanol should be evaluated for use in traps for saproxylic insects as well as their predators and associates. The purchase of isopropanol is unregulated and it could serve as an all-in-one attractant, killing agent, and preservative. Isopropanol thus reduces labor and expenses both at the point of purchase and by eliminating the need for slow-release preparations and mechanisms.

In decomposition processes, isopropanol is either an intermediary or minor end product, and thus may not be as attractive as the major end products such as ethanol. This limitation might also be an important advantage: isopropanol-baited traps could still indicate some species’ presence and patterns of abundance, while making it less likely traps will be overwhelmed by the rapid capture of large numbers of insects.

## ACKNOWLEDGMENTS

Campbell County Public Schools, Lynchburg City Schools Education Foundation, and the ECG Foundation provided financial support for this work. We are greatly indebted to the Pickering, Mabery, Rivers, and Styrsky families for assistance and property access. Mark Deyrup, John Lepri, Katrina Pickering, and Erin Rierson kindly provided valuable feedback that substantially improved the manuscript.

## DATA AVAILABILITY

The data in this study have been deposited in the Harvard Dataverse.

DOI: https://doi.org/10.7910/DVN/HIXSCX

URL: https://dataverse.harvard.edu/api/access/datafile/7568647

